## Supplement Table 1 for "The protective roles of Eugenol on type 1 diabetes mellitus through NRF2 mediated oxidative stress pathway"

**Table 1. Antibodies**

| **Antibody** | **Species** | **Company (catalogue)** | **Dilution** | |
| --- | --- | --- | --- | --- |
|  |  |  | **WB** | **IHC/IF** |
| Insulin | Rabbit | Abcam (ab181547) | 1: 1000 | 1: 200 |
| γH2AX | Rabbit | Abmart (ab208670) | 1: 5000 | 1: 100 |
| BCL2 | Mouse | Affinity (BF9103) | 1: 1000 | ND |
| BAX | Mouse | [Santa Cruz Biotechnology](http://www.baidu.com/link?url=JyKcE01MHDo2C82_vA7PLGsOo6FHhidzFLsl0vS1U__) (sc-7480) | 1: 500 | ND |
| Cleaved Caspase-3 | Rabbit | Cell Signaling Technology (#9664) | 1: 1000 | ND |
| NRF2 | Rabbit | Proteintech (16396-1-AP) | 1: 1000 | 1: 200 |
| KEAP1 | Mouse | Proteintech (60027-1-Ig) | 1: 1000 | ND |
| HO-1 | Rabbit | Proteintech (10701-1-AP) | 1: 1000 | 1: 200 |
| NQO-1 | Mouse | Proteintech (67240-1-Ig) | 1: 1000 | ND |
| β-actin | Rabbit | Proteintech (20536-1-AP) | 1: 5000 | ND |
| LaminB | Rabbit | Proteintech (12987-1-AP) | 1: 1000 | ND |
| GAPDH | Rabbit | Affinity (AF7021) | 1: 5000 | ND |

ND = Not detected; WB = Western blot; IHC: Immunohistochemistry; IF:Immunofluorescence.
