## Supplement Table 2 for "The protective roles of Eugenol on type 1 diabetes mellitus through NRF2 mediated oxidative stress pathway"

**Table 2. Primer information for mouse**

| **Gene name** | **Primer direction** | **Sequences (5’to 3’)** |
| --- | --- | --- |
| *Ins1* | Forward | CAAACCCACCCAGGCTTTTG |
|  | Reverse | AACGCCAAGGTCTGAAGGTC |
| *Bax* | Forward | ACACTGGACTTCCTCCGTGA |
|  | Reverse | AGAGGAGGCCTTCCCAGC |
| *Bcl2* | Forward | TGAACTGGGGGAGGATTGTG |
|  | Reverse | CAGAGACAGCCAGGAGAAATCA |
| *Nrf2* | Forward | CAGCCATGACTGATTTAAGCAG |
|  | Reverse | CAGCTGCTTGTTTTCGGTATTA |
| *Ho-1* | Forward | TCCTTGTACCATATCTACACGG |
|  | Reverse | GAGACGCTTTACATAGTGCTGT |
| *β-actin* | Forward | CTACCTCATGAAGATCCTGACC |
|  | Reverse | CACAGCTTCTCTTTGATGTCAC |
| *Keap1* | Forward | GACTGGGTCAAATACGACTGC |
|  | Reverse | GAATATCTGCACCAGGTAGTCC |
| *Nqo-1* | Forward | GAAGACATCATTCAACTACGCC |
|  | Reverse | GAGATGACTCGGAAGGATACTG |
